# Temporal Self-Compression: Behavioral and neural evidence that past and future selves are compressed as they move away from the present

**DOI:** 10.1101/2021.01.22.427831

**Authors:** Sasha Brietzke, Meghan L. Meyer

**Author notes:** Classification: Psychological and Cognitive Sciences.

## Abstract

Although it is well-known that people feel disconnected from their past and future selves, the underlying mechanism supporting this phenomenon is unknown. To help fill this gap, we considered a basic principle of perception. As objects increase in distance from an observer, they also become logarithmically compressed in perception (i.e., not differentiated from one another), making them hard to distinguish. Here, we report four studies that suggest we may feel disconnected from distant selves, in part, because they are increasingly indiscriminable with temporal distance from the present self. In Studies 1-3, participants made trait ratings across various time points in the past and future. We found that participants compressed their past and future selves, relative to their present self. This effect was preferential to the self and could not be explained by the alternative possibility that individuals simply perceive arbitrary self-change with time irrespective of temporal distance. In Study 4, we tested for neural evidence of temporal self-compression by having participants complete trait ratings across time points while undergoing functional magnetic resonance imaging (fMRI). Representational similarity analysis (RSA) was used to determine if neural self-representations are compressed with temporal distance, as well. We found evidence of temporal self-compression in areas of the default network, including medial prefrontal cortex (MPFC) and posterior cingulate cortex (PCC). Specifically, neural pattern similarity between self-representations was logarithmically compressed with temporal distance. Taken together, these findings reveal a “temporal self-compression” effect, with temporal selves becoming increasingly indiscriminable with distance from the present.

**Significance Statement:** For centuries, great thinkers have struggled to understand why we feel disconnected from our past and future selves. Insight may come from a basic principle of perception: as objects become distant, they also become less discriminable, or ‘compressed.’ In Studies 1-3, we demonstrate that people’s ratings of their own personality become increasingly less differentiated as they consider more distant past and future selves. In Study 4, we found neural evidence that the brain compresses self-representations with time, as well. When we peer out a window, objects close to us are in clear view whereas distant objects are hard to tell apart. We provide novel evidence that self-perception may operate similarly, with the nuance of distant selves increasingly harder to perceive.

## Introduction

Despite inhabiting the same body throughout their lives, humans are remarkably disconnected from their past and future selves. People distance themselves from their past self by misconstruing and disparaging their former qualities (Quoidbach et al., 2013; A. E. Wilson & Ross, 2001), which compromises self-compassion and personal growth (Neff & Pommier, 2013; Zhang & Chen, 2016). On the flipside, people overestimate their future self-improvement and mispredict their future emotions (Armor & Taylor, 2002; Regan et al., 1995; Robins & Beer, 2001; Weinstein, 1980; T. D. Wilson & Gilbert, 2005), producing self-sabotaging behaviors such as procrastination (Blouin-Hudon & Pychyl, 2015) and failing to save resources (Hershfield, 2011). While such findings suggest people treat their past and future selves differently from their present self, how self-continuity breaks down remains to be determined.

Insight may come from cognitive psychology and neuroscience research on how distance from an origin point impacts perception and mental representation of later points. Weber-Fechner law refers to the observation that across perceptual domains (vision, hearing, taste, touch, smell), physical changes in stimuli are logarithmically compressed in perception such that the farther they are from an original stimulus, the less well people differentiate between them (Fechner, 1948). Specifically, in the cognitive sciences, ‘compressed representation’ refers to the phenomenon in which representations do not show the same degree of acuity for all parts of the scale on which they are measured, with later ends of the scale harder to tell apart (i.e., ‘compressed’) than earlier ends of the scale (Fechner 1948; Howard, 2018).

Representations outside of direct perception demonstrate a similar compression phenomenon. The internal representation of numbers abides by logarithmic compression, including in the neuronal code of monkey prefrontal cortex (Dehaene, 2003; Nieder et al., 2002; Shepard et al., 1975). For example, in humans, numbers of greater magnitude become less distinguishable on the number line than numbers of smaller magnitude (Longo & Lourenco, 2007; Dehaene & Mehler, 1992; Piazza, Izard, Pinel, LeBihan, & Dehaene, 2004; Howard, 2018). Moreover, there is evidence that memories are also logarithmically compressed with time: the farther from the present the memory, the less discriminable it is from an earlier memory (Arnold et al., 2016; Jeunehomme et al., 2018; Michelmann et al., 2019). Indeed, it has been suggested that it is easier to recall recent events because they “pop out” during retrieval whereas it is easier to knit distal memories together because the far past has reduced representational acuity between memories (Howard, 2018).

Given that information is compressed with distance across several psychological domains, might the same principle apply to self-representation? Although to our knowledge no research has tested whether self-representations are temporally compressed (i.e., logarithmically compressed with temporal distance from the present), there is social psychological evidence consistent with this possibility. People think their distant future self will be less nuanced than their more recent and present self (Wakslak et al. 2008; Nussbaum, Trope, & Liberman, 2003; Pronin & Ross, 2006), which suggests a lack of uniqueness in distant temporal self-representations. There is also extensive evidence in support of construal level theory from social psychology (Soderberg et al., 2015), which contends that when we reason about distal pasts and futures (including distal past and future *selves*), we focus on more abstract (vs. concrete) features (Trope & Liberman, 2010). Thus, construal level theory is compatible with the idea of logarithmic compression, as temporally compressed past and future selves would provide less discriminable features to reason about, and hence facilitate relying on abstractions. Despite these clues, to date, it remains unknown whether distant past and future selves, like other perceptions and cognitions, are compressed with distance from the present self.

The current research bridges social psychological findings on changes in self-perception across time with psychophysical principles regarding compressed representations to gain traction on how self-continuity may break down as people reflect on themselves farther out in time. Our first goal was to assess the possibility of *‘temporal self-compression’*: whether changes in self-perception are logarithmically compressed such that the farther they are from the present, the less differentiated they become. In Studies 1-3, we leveraged social psychological findings that people tend to perceive they will be better versions of themselves in the future (Armor & Taylor, 2002; Regan et al., 1995; Weinstein, 1980) and were worse versions of themselves in the past (Quoidbach et al., 2013; A. E. Wilson & Ross, 2001) to test our predictions. Participants rated themselves on multiple positive personality traits across various points in their past and future. The temporal self-compression hypothesis posits that the amount of perceived future self-improvement (e.g., in confidence) will be greater between time points closer to the present than between time points farther in the past and the future. Just as numbers of greater magnitude become less distinguishable on the number line than numbers of smaller magnitude, selves of greater distance from the present self will become less distinguishable in their amount of change than selves closer to the present self.

Our second goal was to seek neural evidence of temporal self-compression. Assessing this possibility is critical to the temporal self-compression hypothesis, as it would suggest that self-report evidence of self-compression is not wholly due to biased reporting, but rather may reflect an underlying self-representation that is compressed with time. There is extensive evidence that self-representations are supported by the brain’s default network (Buckner et al., 2008; Northoff et al., 2006), particularly medial prefrontal cortex (MPFC; Denny et al., 2012; Lieberman et al., 2019). For example, the MPFC has been robustly implicated in self-reflection tasks, such as assessing one’s own personality traits (Fossati et al., 2003; Kircher et al., 2002; Ochsner et al., 2005), reflecting on their own emotions (Gusnard et al., 2001; Ochsner et al., 2004), and engaging in self-affirmation (Cascio et al., 2016; Cooper et al., 2015; Falk et al., 2015). To date, brain imaging research investigating the impact of time points on self-representation has found that MPFC is more associated with reflecting on the present self versus the past and future self (D’Argembeau et al. 2008; Tamir and Mitchell 2011; Ersner-Hershfield et al. 2009). However, no brain imaging research has parameterized the distance into the past and future with which participants consider the self, and thus it is unknown whether self-representation in the MPFC follows a temporal self-compression principle.

To determine whether brain regions, particularly MPFC, show evidence of temporal self-compression, in Study 4, participants considered their past and future selves’ positive and negative personality traits across multiple time points while undergoing functional magnetic resonance imaging (fMRI). Our brain imaging paradigm further allowed us to arbitrate between two possible ways in which the brain may compress self-representations with time. One possibility is that past and future selves are compressed as their own, separate representations: past selves may be represented separately in the brain from future selves, distinctly compressed away from the present self. Because the brain imaging paradigm included positive and negative traits, we were able to test whether above and beyond valence differences between appraisals of past and future selves (Wilson & Ross, 2001; Armor & Taylor, 2002; Regan et al., 1995; Weinstein, 1980), they may be distinctly represented from one another and separately compressed away from the present. The alternative possibility, however, is that the temporal compression of self-representations is organized irrespective of past versus future. That is, past and future selves may be collectively compressed with time away from the present. This possibility is consistent with cognitive neuroscience research suggesting that prospection and retrospection rely on overlapping neural mechanisms in the default network, including MPFC (Addis et al., 2009; Schacter et al., 2007). Multivariate pattern similarity analysis was used to determine which of these competing possibilities underlie temporal self-compression. Across studies, if findings support our hypotheses, they would provide the first evidence of temporal self-compression and how it is reflected in the brain.

## Results

### Study 1

The goal of Study 1 was to test whether perceptions of past and future selves are logarithmically compressed with distance from the present. Participants (N=178) rated themselves on 20 positive personality traits (0 not all - 100 extremely; see Figure 1a). Positive personality traits were selected to be in line with past social psychology research, which consistently shows that individuals appraise their past self less positively than their present self and their future self more positively than their present self (Armor & Taylor, 2002; Regan et al., 1995; Weinstein, 1980; A. E. Wilson & Ross, 2001), while still testing the novel prediction of temporal self-compression. Personality traits for all studies were normed for perceived change using ratings from an independent sample (see Methods), and controlled for likeability using the Dumas person-descriptive word list (Dumas et al., 2002). Each trait was rated across 9 time points, up to 1 year in the future and 1 year in the past. The 1 year range was selected because past work shows that considering the self as early as 1 year away from the present is enough to show disconnection from the self (Tamir & Mitchell, 2011). This observation allowed us to therefore ask whether self-perceptions are compressed as they move closer to this distance. Every time scale rated was spaced 3 months apart. Trait presentation was fully randomized across time points.

**Figure 1.**
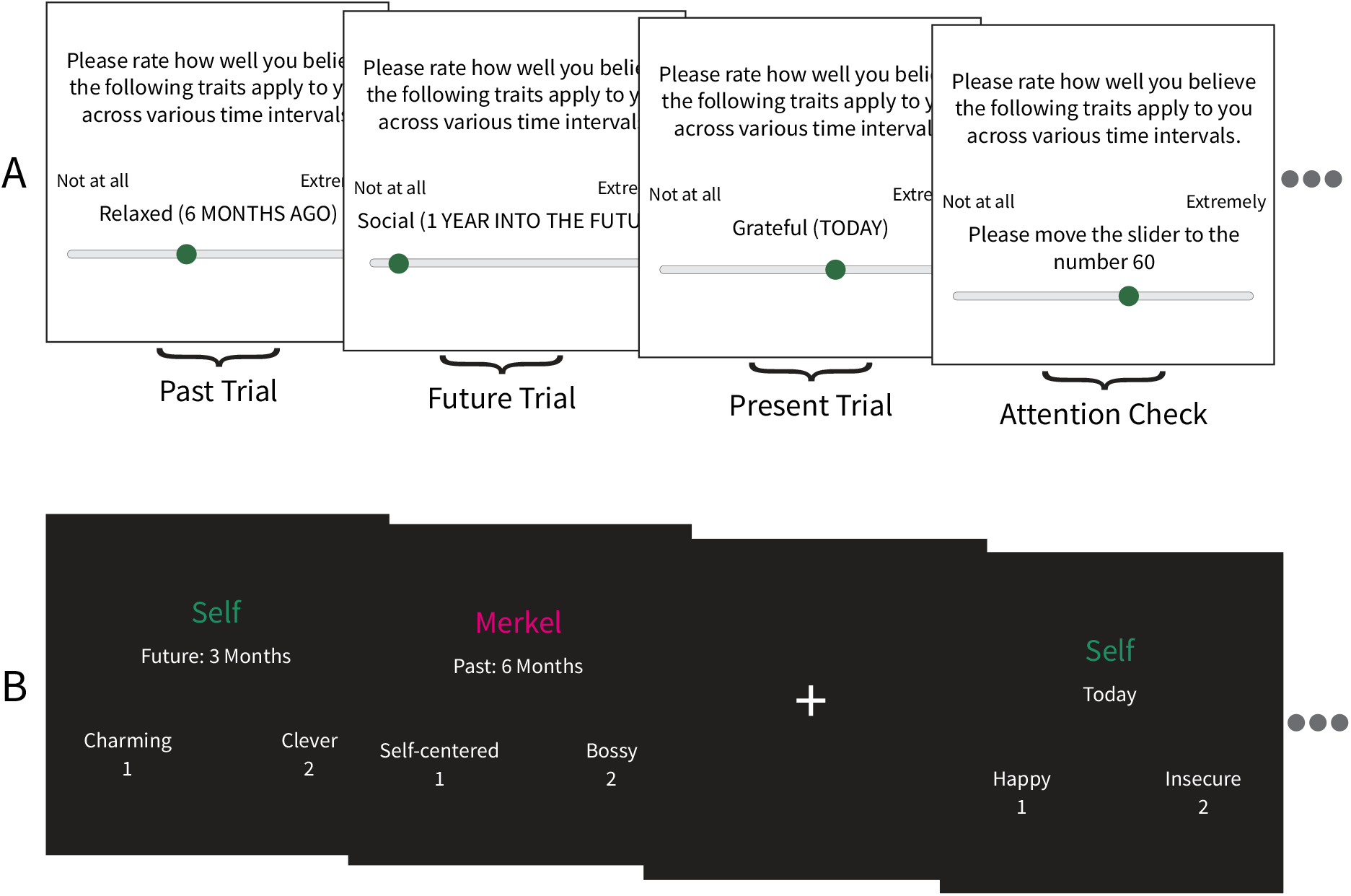
Paradigm schematics. (A) In the behavioral paradigm for Studies 1-3, participants rated themselves on positive personality traits on a sliding scale (0 not all - 100 extremely) across 9 time points in the past and future. Attention checks that prompted participants to move the slider to a certain value were included to assess data quality. (B) Across 4 runs in the fMRI paradigm, we asked participants to choose which of two traits either self or, separately, Merkel embodied more across 9 time points in the past and future. Each trait was presented for 5 seconds and included 30% jittered trials of fixation.

A linear mixed model revealed a significant linear (β=2.25, standardized β=0.24, t-statistic=10.14, df=82.78, p<0.001) and cubic (β=−2.17, standardized β=−0.09, t-statistic=−8.36, df=31645.86, p<0.001) relationship of time on personality trait rating. As shown in Figure 2a, there are rapid changes in self-perception occurring in the time periods adjacent to the present, which then tapers out with increased temporal distance. Note that the cubic effect in this instance is consistent with past work finding logarithmic compression with distance because participants are demonstrating logarithmic compression in separate directions into the past and future, with less positive self-perceptions into the past and more positive self-perceptions into the future.

**Figure 2.**
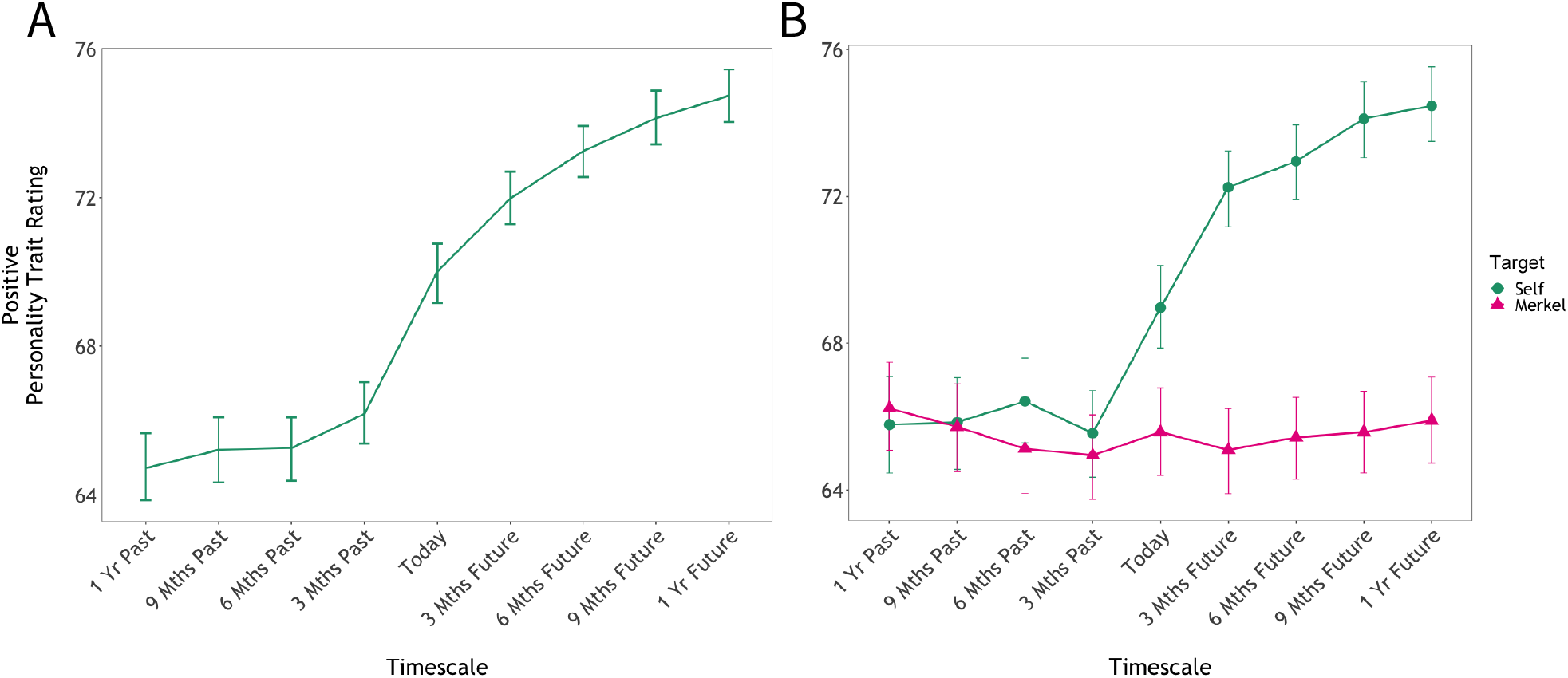
(A) Study 1 demonstrates significant temporal self-compression of personality trait ratings. (B) Study 2 demonstrates that temporal compression is preferential to the self, relative to another well-known person, Angela Merkel.

Given past research showing age impacts temporal self-appraisal (Addis et al., 2008; Levine et al., 2002), we also assessed our model including age as a covariate to rule out the possibility that our findings were solely driven by age. While older people did rate themselves more positively (β=7.87, standardized β=0.09, t-statistic=2.30, df=177.46, p=0.021) and exhibited less linear (β=−2.29, standardized β=−0.07, t-statistic=−3.96 df=390.96, p<0.001) and cubic compression (β=2.27, standardized β=0.03, t-statistic=2.47, df=31645.90, p=0.014) compared to younger adults, our primary effect of cubic compression of trait ratings remained significant when age was included in the model (β=−10.17, standardized β=−0.09, t-statistic=−3.25, df=31645.90, p=0.002).

### Study 2

We next aimed to determine if temporal self-compression is relatively preferential for the self, or whether it extends to other people more generally. Consistent with past research on the self (e.g., Krienen et al., 2010), we selected a politician for the other-person condition: German chancellor Angela Merkel. To ensure all participants had similar knowledge about Angela Merkel, participants were provided a short biography of Merkel prior to performing their ratings. Participants (N=174) rated themselves and Merkel on the same personality traits used in Study 1 and across the same time points (i.e., present, past, and future ratings spaced 3 months apart up to 1 year).

A linear mixed model demonstrated a significant interaction effect between linear time and target (β=2.06, standardized β=0.21, t-statistic=8.94, df=30953.38, p<0.001), and cubic time and target (β=−2.01, standardized β=−0.08, t-statistic=−3.39, df=30953.38, p=0.001), such that the ratings for self displayed stronger linear and cubic effects relative to Merkel. In other words, and as shown in Figure 2b, temporal compression was preferential to self-ratings (vs. Merkel ratings). It is noteworthy that we also observed a significant effect of target on personality trait ratings (β=4.08, standardized β=0.16, t-statistic=17.97, df=30953.38, p<0.001), such that ratings for the self were higher than ratings for Merkel.

Although there was no significant effect of age on trait ratings when both targets were considered simultaneously (β=1.00, standardized β=0.01, t-statistic=0.29, df=178.06, p=0.775), there was a significant interaction between age and target (β=11.06, standardized β=0.13, t-statistic=13.94, df=30953.37, p<0.001), such that trait ratings were higher among older adults for self, but not for Merkel. Consistent with Study 1, even after controlling for age (i.e., adding it as a covariate), there was a significant interaction between linear time and target (β=2.06, standardized β=0.21, t-statistic=8.96, df=30953.37, p<0.001), and cubic time and target (β=−2.01, standardized β=−0.08, t-statistic=−3.40, df=30953.37, p=0.001), suggesting age-related effects do not explain temporal self-compression.

### Study 3

Results from Studies 1-2 are consistent with the temporal self-compression hypothesis. However, an alternative interpretation of these results is that individuals are accessing a generic rating of their perceived self-change invariant to the time point considered. People may have a theory of self-change that is logarithmic regardless of the actual time point into the past or future considered. In other words, temporally distant selves may not be compressed, but rather represent a categorical difference in self-perception from the present self. To rule out this possibility of “arbitrary self-improvement”, we would need to include time points closer to the present than 3 months. If changes in self-perceptions are compressed specifically as people consider more distant time points, as we propose in the temporal self-compression hypothesis, then the compression should *not* occur for times closer to the present. If this were the case, then we would expect two findings: 1) earlier temporal time points (e.g., the self in 1 week or 1 month) should not be rated equivalently to more temporally distant time points (i.e., 3 months and beyond) and 2) the amount of perceived self-change in smaller increments of time (e.g., 1 day vs. one week) should be *similar* to the perceived amount of change in later, longer intervals of time (e.g., 9 months vs. 1 year). This latter point would reflect that the amount of self-change one can imagine in their more distant future is compressed relative to the amount of change during earlier, smaller time intervals. It also fits with the idea of ‘scale-invariance’ in logarithmically compressed representations (Howard, 2018). For example, a consequence of the observation that numbers of greater magnitude become less distinguishable on the number line than numbers of smaller magnitude is that the difference between, for example 10 and 11, is perceived as more similar to the difference between 100 and 110 than to the difference between 100 and 101 (Longo & Lourenco, 2007; Dehaene & Mehler, 1992; Piazza, Izard, Pinel, LeBihan, & Dehaene, 2004; Howard, 2018; see Figure 3a for a schematic of the temporal self-compression hypothesis vs. the alternative, arbitrary self-improvement hypothesis). Likewise, the amount of perceived change in the self between tomorrow and one-week may counterintuitively be similar to the amount of perceived change in the self between 3 months and 6 months, despite the latter capturing a longer interval of time.

**Figure 3.**
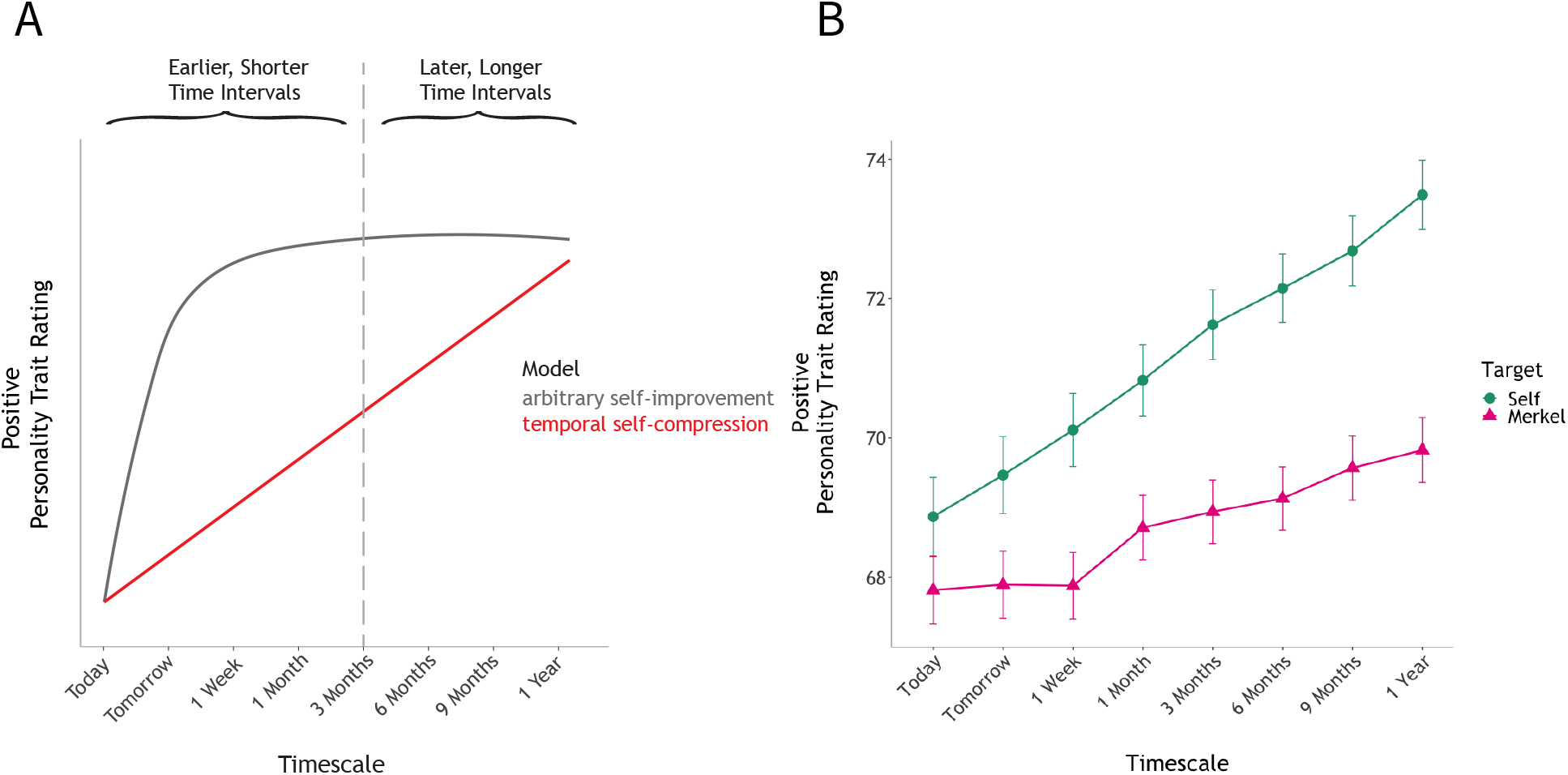
Results for Study 3. (A) Schematic of two competing possibilities for self-perception: temporal self compression (red line) vs. arbitrary self-improvement (grey line). The “arbitrary self-improvement” model demonstrates the possibility that individuals access a generic rating of future self-improvement. The “temporal self-compression” model predicts a gradual, not abrupt, transition when earlier time points with smaller intervals of time are assessed. (B) Empirical findings abide by the temporal self-compression model: Study 3 demonstrates a significant linear trend for self-ratings that is preferentially stronger for the self (vs. a well-known other, Angela Merkel).

To adjudicate between the competing possibilities of temporal self-compression vs. arbitrary self-improvement, in Study 3, participants (N=192) made their personality trait ratings for the self and Merkel using the following intervals into the future: today, tomorrow, 1 week, 1 month, 3 months, 6 months, 9 months, and 1 year. We focused on the future specifically given that Studies 1 and 2 showed effects for both past and future (and thus it was not necessary to include both here) and to minimize participant burden given the additional temporal conditions.

Consistent with the temporal self-compression prediction laid out above, a linear mixed model revealed a significant interaction between time and target (β=0.34, standardized β=0.04, t-statistic=3.95, df=30324.0, p<0.001), demonstrating the linear effect of perceived self-improvement was stronger for the self than for Merkel (see Figure 3b). In other words and as shown in Figure 3b, we saw evidence of strong linear self-improvement with time demonstrating that 1) earlier time points are not rated equivalently to later time points and 2) the rate of perceived self-change between smaller temporal increments at earlier time points is similar to the rate of perceived self-change between larger, more distant time points. There was also a significant effect of time (β=0.31, standardized β=0.03, t-statistic=3.10, df=288.1, p=0.002) and target (β=0.88, standardized β=0.11, t-statistic=2.01, df=30324.0, p<0.044), such that personality trait ratings were higher for self and for time points farther ahead in the future. Critically, if the arbitrary self-improvement account was correct, then when compression terms are added to the model, the interaction between logarithmic time (used instead of a cubic term because this study only included the future) and target would be statistically significant, but that was not the case (compression term β=0.27, standardized β=0.01, t-statistic=0.38, df=30324.0 p=0.701). Thus, the arbitrary self-improvement account can be ruled out.

We also assessed our model including age as a covariate. Replicating our previous results, we found older people rated themselves more positively (β=15.18, standardized β=0.20, t-statistic=4.50, df=191.74, p<0.001) and exhibited less of a linear trend across time (β=−1.15, standardized β=−0.02, t-statistic=−3.78, df=192.0, p<0.001) compared to younger adults. Critically, the interaction between logarithmic time and target was still not statistically significant (β=−0.17, standardized β=0.01, t-statistic=0.38, df=30324.02, p=0.701) when age was controlled for (i.e., added as a covariate).

### Study 4

The goal of Study 4 was to search for neural evidence of temporal self-compression. This would not only add additional support for the hypothesis; it would further suggest that results from Studies 1-3 are not merely an artifact of self-report bias (Nisbett & Wilson, 1977), but rather may reflect an underlying mental representation of the self that is compressed with time. If self-representations are temporally compressed, then we would expect brain regions associated with self-representation to show less neural pattern similarity between the present self and selves 3 months away and greater neural pattern similarity between selves as participants reflect farther out in time (e.g., neural pattern similarity between 3 months away and 6 months away; 6 months away and 9 months away; 9 months away and 12 months away). Moreover, we assessed which of two competing possibilities best explain how the brain temporally compresses past and future selves: whether past and future selves are compressed as their own, separate representations or whether past and future selves are represented similarly to one another and collectively compressed with time away from the present.

Participants in Study 4 (N=43) completed a trait reflection task while undergoing functional magnetic resonance imaging (fMRI). For the scanner task, participants chose which of two traits best reflected themselves (self condition) separately across the 9 time scales used in Study 1 and 2 (i.e. Future-3 Months). Participants also completed trait reflection trials across these time points for Merkel. As in our previous studies, participants received a short biography of Angela Merkel before completing the task. Critically, in Study 4, one-third of trials showed two positive trait options (e.g., charming; wise), two negative trait options (e.g., insecure; lazy), or a combination of positive and negative traits (e.g., charming; insecure). Including positive and negative traits helps ensure that neural results reflect a compressed self-representation as opposed to changes in similarity in valence more generally. As is the case for the positive traits, negative traits were normed on perceived change and controlled for likeability using the Dumas word list. Every subject completed the same number of positive-positive, negative-negative, and positive-negative trait rating trials for their scans although trait pairings were counterbalanced between self and Merkel targets to help ensure participant engagement.

To test the competing possibilities of temporal self-compression, we employed representational similarity analysis (RSA; Kriegeskorte et al. 2008) to search for regions across the brain that may fit the competing predictions. We constructed two theoretical representational dissimilarity matrices (RDMs): a sigmoidal model differentiating past and future selves, and a logarithmic model in which past and future selves are represented similarly (see Figure 4a). Both models assume temporal self-compression (i.e., greater similarity in representations further out in time with dramatic shifts closer to the present) and only differ in whether past and future selves are separately vs. collectively represented.

**Figure 4.**
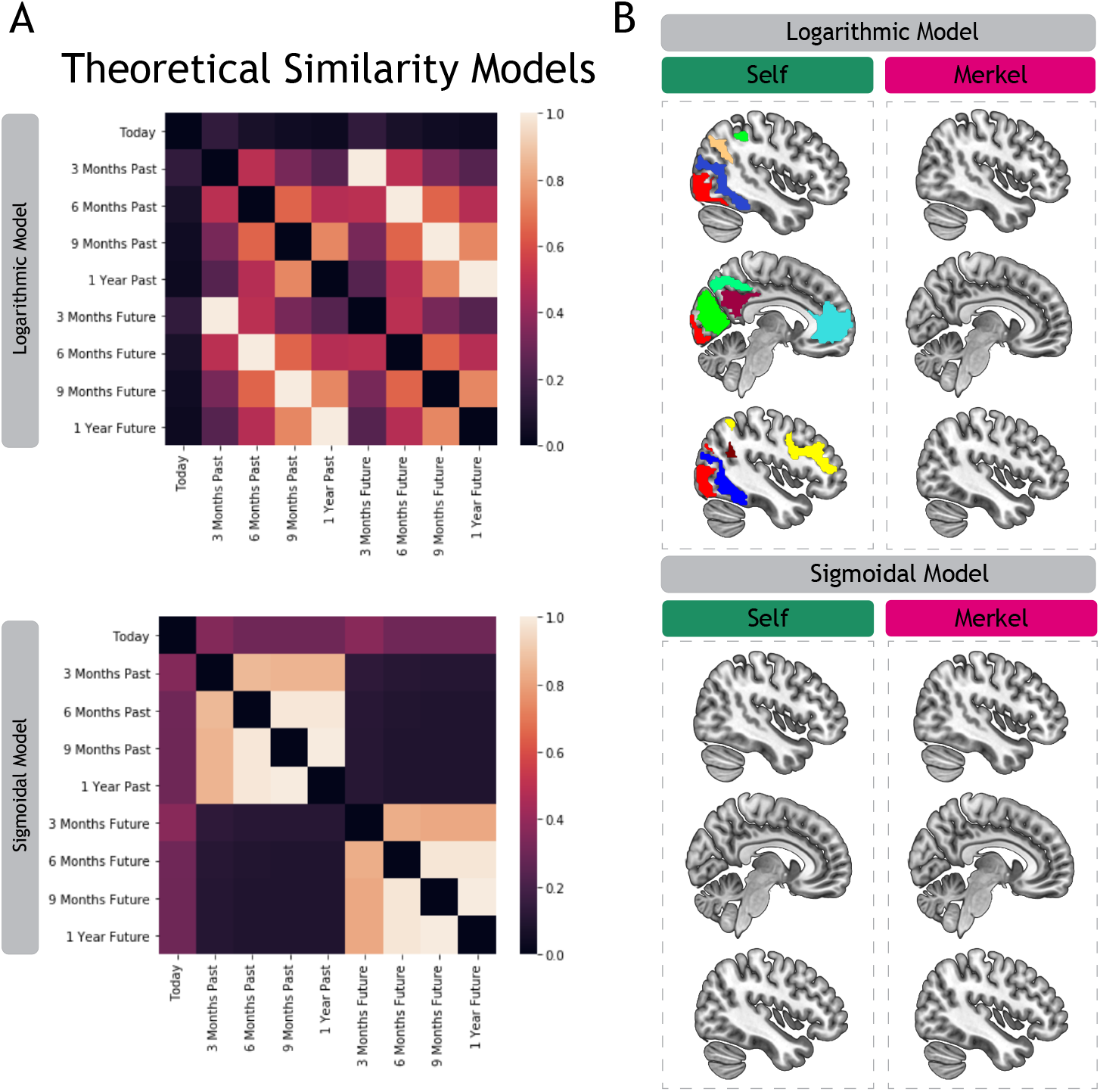
Whole brain representational similarity analysis (A) We constructed two theoretical representational dissimilarity matrices (RDMs): a logarithmic model in which past and future selves are represented similarly and a sigmoidal model differentiating past and future selves. Each RDM is symmetrical about a diagonal of zeros and only the vectorized lower triangle is extracted (without the diagonal) (Kriegeskorte et al., 2008), (B) Spearman rank correlations were conducted between each RDM and each region of interest in the Yeo parcellation scheme, for self and Merkel beta images separately. Brain regions across the brain, including default network regions previously associated with self-reflection (e.g., MPFC; PCC), were significantly associated with the logarithmic model for the self (FDR < 0.05). No brain regions were significantly associated with any of the other models.

We looked for significant correlations between the two theoretical structures across targets (self and Merkel) across the whole-brain. The sigmoidal model, which would reflect past and future selves are separately and distinctly represented and temporally compressed, yielded no statistically significant results for either the self or Merkel (see Figure 4b). However, the logarithmic model, which reflects past and future selves as collectively represented and compressed with time, showed significant results in brain regions previously associated with self-representation: MPFC, as well as other default network regions (posterior cingulate cortex (PCC), bilateral precuneus, right middle temporal gyrus, (for full list of regions see Supplementary Table 1; p<0.05, FDR-corrected for multiple comparisons). Additional regions also showed this relationship, including regions in the frontoparietal control network, dorsal attention network, and visual system (see Supplementary Table 1). In contrast to the self condition, no regions of the brain were associated with the logarithmic model for the Merkel condition (see Figure 4b).

### MPFC Region-of-Interest (ROI) Pattern Similarity Analysis

We next sought to further probe whether MPFC voxels in the brain specifically associated with self-representation adhere to the logarithmic model. To assess MPFC voxels associated with self-representation defined independently of our own data, we applied a Neurosynth-derived MPFC ROI using the search term ‘self’; this ROI has been used in past research on the self (Meyer & Lieberman, 2018; Courtney and Meyer 2020). For every contiguous time point (separately for the self and other; e.g., similarity between Self_Today_ and Self_3-MonthsFuture_), similarity between the two MPFC ROI beta images was calculated via Spearman rank correlations within each subject (see Figure 5a for a schematic of the analysis plan).

**Figure 5.**
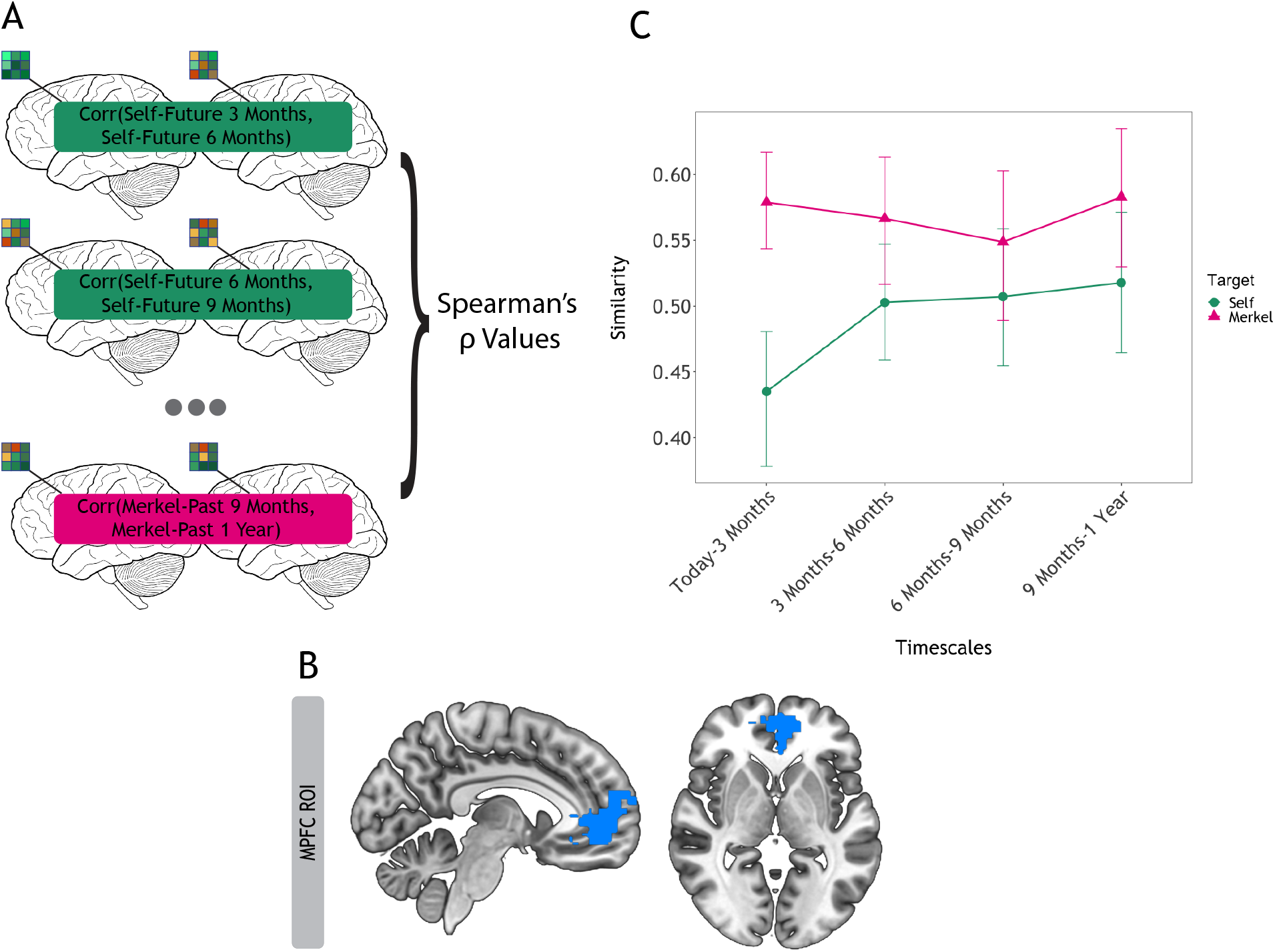
Self-specific MPFC Similarity Analysis. (A) Schematic of the analytic procedure. For every contiguous time point for both the self and other (e.g., Self-Today and Self-3 Months in the Future), neural pattern similarity between the two masked beta images was calculated via Spearman rank correlations within each subject. (B) Neurosynth-derived MPFC ROI using the search term ‘self’ used in this analysis. Voxels in the association map test were restricted to Brodmann Area 10, a region particularly associated with self-representation. (C) Results show a significant interaction effect between logarithmic time and target, meaning that the MPFC pattern similarity displayed between contiguous time points across time (today to 1 year) was preferentially logarithmically compressed with time for the self (vs. another well-known person, Angela Merkel).

A linear mixed model revealed a significant interaction effect between logarithmic time and target (β=0.06, standardized β=0.15, t-statistic=2.50, df=546, p=0.013), meaning that the MPFC pattern similarity displayed between contiguous time points across time (today to 1 year) was more logarithmic for the self than for Merkel (see Figure 5c). Put simply, neural pattern similarity for the self in the MPFC was lowest for the most proximal time points (i.e., more differentiated), then increased in similarity when comparing more distal time points (i.e., less differentiated (β=0.06, standardized β=0.14, t-statistic=2.47, df=39, p=0.013). The same relationship was not statistically significant when considering Merkel (β=−0.01, standardized β=− 0.01, t-statistic=−0.31, df=273, p=0.75). The linear mixed model also identified an effect of target on similarity value across time (β= −0.13, standardized β=−0.35, t-statistic=−5.30, df=546, p<0.001), such that MPFC similarity between contiguous time points (e.g., Self_Today_ and Self_3-MonthsFuture_) were lower for the self than for Merkel.

## Discussion

Across domains studied outside of social psychology, changes in stimuli are logarithmically compressed such that the farther they are from an original stimulus, the less discriminable they become. In four studies, we found novel evidence suggesting temporal self-perception abides by this principal as well. Study 1 demonstrates that people compress their future and past selves such that distal selves are perceived more similarly to one another compared to current and proximal selves. Study 2 replicates this finding and further shows that this compression effect may be relatively preferential to the self. Consistent with Weber Fechner’s law that *distant* perceptions specifically are compressed, Study 3 showed that temporal self-compression occurs at more distant but not more proximal time points. Finally, temporal self-compression is not simply due to a bias in self-report, given that in Study 4, we found that neural representations of past and future selves are also collectively compressed with time away from the present.

Our findings may help explain some of the counterintuitive ways people treat their temporally distant selves. For example, an extensive literature on hyperbolic discounting suggests that people display preferences for selecting a smaller monetary reward immediately rather than waiting to receive a larger sum (Ainslie, 1975). Because this formulation is hyperbolic, rather than exponential, the value of *immediate* reward is unduly high such that people succumb to instant gratification; yet, when asked to choose between two temporally distant choices, people manage to exhibit more restraint. Why would multiple future selves be short-changed for the present self in a time-inconsistent way? Our results indicate that more distant future selves are not easily distinguishable from one another. The representational similarity between more distant selves would thus make it easier to not treat them differently from one another and hence be willing to wait for the larger reward—it’s the “same” self anyways. Relatedly, people are less prone to hyperbolic discounting when making decisions for *others* (Albrecht et al. 2011) and recent research has shown that representations of others’ internal states (in the present) are also less differentiated from one another than internal representations of the self’s states (in the present) (Thornton et al., 2019). This further speaks to the possibility that difficulty distinguishing between distant self-representations may facilitate delayed gratification in choices.

We replicate past social psychology research demonstrating people perceive their past selves less positively than their present self and their future self as even better (Armor & Taylor, 2002; Regan et al., 1995; Weinstein, 1980). This phenomenon is thought to occur for motivational reasons: people derogate their past selves to boost their current self-worth (A. E. Wilson & Ross, 2001) and see their future self through rose colored glasses to sustain optimism for their future (Taylor, 1989). Our novel addition to this literature is that this bias in self-perception follows basic principles of other forms of perception and mental representation: as individuals consider more distant past and future selves, their ability to perceive them as uniquely worse or better is increasingly attenuated.

Our findings, in conjunction with the prior literature on positive self-illusions (Robins & Beer, 2001; Taylor, 1989), help explain why we did not observe temporal compression for the non-self target (i.e., Merkel). Without the motivation to treat a person differently in the past and future, perceptions of that person have no direction to move away from the present, which provides no opportunity for compression with distance. This suggests that we may temporally compress representations of other people we know but are also motivated to treat differently over time. For example, temporal perceptions of friends and enemies may likewise be compressed with time, possibly in opposing directions in valence. Similarly, individual differences in self-perception may moderate the extent and valence of temporal self-compression. For example, individuals with depression, who often perceive the self negatively, may show different rates of temporal self-compression or perhaps even show a present self that is actually part of a negative, compressed past self. The construct of temporal self-compression thus provides novel ideas regarding possible mechanisms to investigate in individuals struggling to see themselves positively. Future research will help determine the extent to which temporal compression is strongest for the self (vs. other individuals we are motivated to see change over time), as well as whether it fundamentally varies for the self across conditions characterized by negative self-views.

Our results also revise the view that the self is ‘special.’ Multiple pieces of evidence point to the possibility that self-representation is unique from other types of knowledge representation (Denny et al., 2012; Sui et al., 2015; Sui & Humphreys, 2015; Symons & Johnson, 1997). For example, self-reflection is preferentially associated with portions of MPFC (Sui and Humphreys 2015; Sui et al. 2015; Denny et al. 2012; Symons and Johnson 1997; Lieberman et al., 2019) and self-relevant information is privileged in memory (Sui et al. 2015; Symons and Johnson 1997). Yet, here we found that self-perception is susceptible to Weber Fechner’s law much like other forms of perception and mental representation. In fact, our MPFC ROI is constrained to brain voxels preferentially responsive to the self, and these voxels demonstrated this ubiquitous principle of compression. How can our results be compatible with past findings suggesting the uniqueness of self-knowledge? One possibility is that self-knowledge may be stored separately from other representations, but the organization of that knowledge abides domain-general principles, such as temporal compression. Future research that includes temporal self-appraisal stimuli along with perceptual stimuli that vary in distance from an origin point can help test this possibility. This approach may also clarify whether regions outside of the default network that showed evidence of temporal self-compression (e.g., portions of the frontoparietal network) may play a domain-general role in this phenomenon.

The present work complements and extends prior neuroimaging findings suggesting that prospection and retrospection rely on the same brain regions. Regions of the default network— not only MPFC, but also precuneus, and medial temporal lobe—are consistently recruited when simulating episodes in the past and thinking about a hypothetical situation in the future (Addis et al., 2007; Okuda et al., 2003; Schacter et al., 2007). Though our behavioral data could suggest past and future selves are represented separately from one another (because past selves were rated as “worse” than future selves), once trial types sampled traits that varied in valence, past and future selves showed no evidence of discrete representation (i.e., the sigmoidal model that searched for past and futures selves separately compressed from the present correlated with no regions in the brain). Instead, we found remarkable overlap between the brain regions that fit the logarithmic model indicating *collective* representation of past and future selves compressed with time and the brain regions that are commonly recruited for prospection and retrospection (Schacter et al., 2007). Such similarities are in spite of the fact that our paradigm tests semantic memory (i.e., personality traits), rather than episodic memory. Our findings are thus compatible with prior work on retrospection and prospection, with the novel addition that temporal selves are logarithmically compressed with time.

Not all age groups compress equally, however. In Studies 1-3, we found that older individuals (vs. younger individuals) displayed less change in self-perception (i.e. attenuated linear and cubic trends) when rating their personality traits across time. In other words, compared to younger adults, older adults view themselves as more static across time. Importantly, our temporal self-compression findings remained significant even when controlling for age. Nonetheless, the aging results are interesting in light of other research demonstrating that older adults generate fewer specific details (or more high level construals) during both retrospection and prospection (Addis et al., 2008; Levine et al., 2002). Aging is further associated with changes in default network function (Andrews-Hanna et al., 2007; Damoiseaux et al., 2007; Spreng & Turner, 2013). On the one hand, this lack of perceived change in the self with time could be due to changes in temporal simulation skills with aging. On the other hand, lack of perceived self-change could reflect motivational changes in self-perception with age. For example, people may be motivated to stabilize their self-views with age if they perceive limited avenues for change; thus considering the self one year out into the past and future, as we did in our studies, may not be distant enough to detect strong self-compression effects in older populations. Relatedly, we replicated findings from the affective aging literature suggesting that older adults display a positivity bias (Carstensen & Mikels, 2005; Mather & Carstensen, 2005). In our studies, older adults appraised themselves better compared to younger adults across positive traits. One direction for future research could be to further unpack the representational structure of temporal selves in older adults in the default network, and the extent to which this may have consequences for behavior.

In summary, we provide the first behavioral and neural evidence to suggest that past and future selves are temporally compressed. Just as the details of physical objects farther out in space are difficult to see, distant past and future selves may be similarly blurry. These findings add new insight into questions of self-continuity over time that have perplexed philosophers and psychologists alike for centuries. Indeed, David Hume suggested that when we lack access to perceptions, self-continuity breaks down (Hume, 1739). Consistent with Hume’s account, our results suggest we may struggle to treat our distant past and future selves as part of our own identity, in part, because they are in less clear view.

## Materials & Methods

### Studies 1-3

#### Participants and Procedures

For Studies 1-3, Amazon Mechanical Turk (MTurk) participants completed an online survey in which they were asked to rate themselves on positive personality traits (0 not all - 100 extremely) across various time points. Positive personality traits were selected to be consistent with past social psychology research, which consistently shows that individuals appraise their past self less positively than their present self and their future self more positively than their present self. Trait presentation was fully randomized across time points. Further, we built in attention checks (i.e., prompting participants to move the sliding scale to 60 instead of rating a trait characteristic). All participants across studies were paid $4.00 for their time and effort.

##### Personality Trait Norming

Personality traits were selected from the Dumas person-descriptive word list (Dumas et al., 2002) and normed for perceived change using ratings from an independent MTurk sample. Participants who normed the stimuli (N=62, 37% Female, mean age=34.8; s.d. 9.6) were instructed to rate 150 positive and negative trait adjectives on a 0-100 scale according to how they perceive the *average person* changing on that trait in one year. For each trait, an average change score was calculated and then z-scored. For Studies 1-3, positive traits were selected such that traits from across the perceived change spectrum were sampled, and controlled for likability using scores from the Dumas word list. In Study 1, 20 traits were selected, whereas in Studies 2-3, 10 traits were selected due to the fact that we were asking participants to rate each trait for the self and Merkel (vs. solely the self in Study 1), and we wanted to ensure participants could complete the tasks in a reasonable amount of time (approximately 30 minutes).

##### Trait Ratings

For Studies 1 and 2, each trait was rated across 9 time points, up to 1 year in the future and 1 year in the past; every time scale was spaced 3 months apart and included the present (see Figure 1a). Studies 2 and 3 prompted participants to rate both the self and German Chancellor Angela Merkel. Study 3 prompted participants to respond across uneven time scales in the future only (today, tomorrow, 1 week, 1 month, 3 months, 6 months, 9 months, 1 year).

##### Participants

In Study 1, 196 participants were recruited from MTurk. In Study 2, 186 participants were recruited from MTurk. In Study 3, 202 participants were recruited from MTurk. For demographic information for all three studies, see Supplementary Table 2. All participants provided informed consent in accordance with the Dartmouth College Institutional Review Board and received $4 for their participation.

#### Data Analysis

After assessing data quality via attention checks, 178 subjects remained in Study 1, 174 subjects remained in Study 2, and 192 subjects remained in Study 3. All code and data for these studies has been made available on OSF (https://osf.io/m458t/). For each study, linear mixed models using the R package *lme4* (Bates, 2007) were constructed to assess how time affected the trait ratings. Linear, quadratic, and cubic terms for time were added as fixed effects to the models in Study 1 and 2. P-values were calculated for mixed model fits by Satterthwaite’s degrees of freedom method. Adding quadratic and cubic terms to the linear-only models in Studies 1-2 significantly improved model fits based on BIC (lowered by 51 and 9 in Study 1 and 2 respectively). In Study 3, we were interested in assessing the arbitrary self-improvement hypothesis, with the hypothesis that when earlier and unevenly spaced time points were assessed, perceived change would be linear (rather than compressed). Thus, the primary model in Study 3 assessed the linear relationship with time. The models all included random intercepts and slopes for participants and random intercepts for item-level traits to account for variability within the population and among traits selected and better ensure generalizability (Judd et al., 2012).

Because Studies 2-3 included both self and Merkel as target conditions, target was added as a fixed effect for those models. In order to identify a compression effect that is preferential to the self in Study 2, the interactive effect of both 1) linear time and target and 2) cubic time and target were added to the model. For the primary model in Study 3, we first added solely the interactive effect of linear time and target to see if there was a linear trend on a compressed time scale. Then, to confirm closer time points are not compressed, we assessed another model for Study 3 with fixed effects of linear, quadratic, and cubic time, target, and the interaction terms between linear and cubic time and target respectively. This latter model including temporal compression terms did not have a statistically better model fit (p-value=0.8696) compared to the model containing solely a linear time term.

Given that age impacts temporal self-appraisal, we also constructed models for each study adding log-transformed age (to account for skewness in the data) as a fixed effect. For Studies 1-3, we included the interaction between age and time (both linear and cubic for Studies 1-2, and just linear for Study 3) to assess whether participants who varied in age displayed the self-compression effect differently.

### Study 4

#### Participants and Procedures

Forty-three right-handed Dartmouth College undergraduates (women 62.8%, mean age=19.1; s.d.=0.88, racial breakdown: 41.9% white, 39.5% Asian, 9.3% African American, and 9.3% Hispanic) were recruited for the present study and screened for any MRI contraindications (e.g., metal in body, claustrophobia, pregnancy). Participants either received extra credit for a course or cash in exchange for their participation. All participants provided informed consent in accordance with the Dartmouth College Institutional Review Board. Three participants ended their scanning session early due to technical difficulties or scanner-induced claustrophobia. Another subject responded to only 61% of trials, and was therefore removed from analysis, bringing the total analysis to 39 subjects.

In the scanner portion of Study 4, we asked participants to choose which of two traits either self or Merkel embodied more across 9 different time scales (the same time points as in Study 1a and 1c; see Figure 1b). Subjects used a button box in the scanner to select which trait they most identified with at that point in time.

One-third of the scanner task trials showed two positive trait options (e.g., charming; wise), two negative trait options (e.g., insecure; lazy) or a combination of positive and negative traits (e.g., charming; insecure). Including positive and negative traits helps ensure that neural results reflect a compressed self-representation as opposed to changes in similarity in valence more generally. As is the case for the positive traits, negative traits were normed on perceived change and controlled for likeability using the Dumas word list. Every subject had the same number of positive-positive, negative-negative, and positive-negative trials for their scans, although trait pairings were counterbalanced between self and Merkel targets to help ensure participant engagement. Each trait was presented for 5 seconds.

#### fMRI Data Acquisition

Brain imaging took place on a Siemens Prisma 3T scanner. Four functional runs in response to the task were acquired using an EPI gradient-echo sequence (2.5 2.5 2.5 mm voxels, TR = 1000 ms, TE = 30 ms, 2.5 mm slice thickness, FOV = 24 cm, matrix = 96 96, flip angle = 59; simultaneous multi-slice (SMS) = 4). A T2-weighted structural image was acquired coplanar with the functional images (0.9 0.9 0.9 mm voxels, TR = 2300 ms, TE = 2.32 ms, 0.9 mm slice thickness, FOV = 24 cm, matrix = 256 256, flip angle = 8). Sequence optimization was obtained using optseq2 (Dale, 1999) and included 30% jittered trials of fixation for measuring a baseline estimation of neural activity.

#### Brain Imaging Data Preprocessing and Beta Estimates

Results included in this manuscript come from preprocessing performed using *fMRIPrep* 1.4.0 (Esteban, Markiewicz, et al. (2018); Esteban, Blair, et al. (2018); RRID:SCR_016216), which is based on *Nipype* 1.2.0 (Gorgolewski et al. (2011); Gorgolewski et al. (2018); RRID:SCR_002502). Per recommendations from the software developers, we report the exact text generated from the boilerplate below.

For each of the 4 BOLD runs found per subject, the following preprocessing was performed. First, a reference volume and its skull-stripped version were generated using a custom methodology of *fMRIPrep*. The BOLD reference was then co-registered to the T1w reference using bbregister (FreeSurfer) which implements boundary-based registration (Greve & Fischl, 2009). Co-registration was configured with nine degrees of freedom to account for distortions remaining in the BOLD reference. Head-motion parameters with respect to the BOLD reference (transformation matrices, and six corresponding rotation and translation parameters) are estimated before any spatiotemporal filtering using mcflirt (FSL 5.0.9, (Jenkinson et al., 2002). BOLD runs were slice-time corrected using 3dTshift from AFNI 20160207 (Cox & Hyde, 1997), RRID:SCR_005927). The BOLD time-series were resampled into standard space, generating a *preprocessed BOLD run in [‘MNI152NLin2009cAsym’] space*. First, a reference volume and its skull-stripped version were generated using a custom methodology of *fMRIPrep*. Several confounding time-series were calculated based on the *preprocessed BOLD*: framewise displacement (FD), DVARS and three region-wise global signals. FD and DVARS are calculated for each functional run, both using their implementations in *Nipype* (following the definitions by (Power et al., 2014). The three global signals are extracted within the CSF, the WM, and the whole-brain masks. The head-motion estimates calculated in the correction step were also placed within the corresponding confounds file. The confound time series derived from head motion estimates and global signals were expanded with the inclusion of temporal derivatives and quadratic terms for each (Satterthwaite et al., 2013). Frames that exceeded a threshold of 0.5 mm FD or 1.5 standardised DVARS were annotated as motion outliers. All resamplings can be performed with *a single interpolation step* by composing all the pertinent transformations (i.e. head-motion transform matrices, susceptibility distortion correction when available, and co-registrations to anatomical and output spaces). Gridded (volumetric) resamplings were performed using antsApplyTransforms (ANTs), configured with Lanczos interpolation to minimize the smoothing effects of other kernels (Lanczos, 1964).

Images that estimated each time period’s task-conditioned effects for both self and other were calculated by modeling the events of interest convolved with the canonical hemodynamic response function in a general linear model (GLM). This model included nuisance regressors for the six motion parameters (x, y, x directions and roll, pitch, yaw rotations), each motion parameter’s derivative and square of the derivative, linear drift, and run constants. We additionally regressed out TRs in non-steady state and TRs that exhibited spikes of motion found from global signal outliers and outliers derived from frame differencing (each 3 standard deviations).

### Representational Similarity Analysis (Whole brain)

Multivariate analyses were all conducted using Python packages, including nltools 0.3.14 (Chang et al., 2019) and nilearn (Abraham et al., 2014). After constructing these representational structures, for each participant, we conducted Spearman rank correlations between each structure and each region of interest in the Yeo parcellation scheme (Yeo et al., 2011) that exceeded 5 voxels, for self and other (i.e., Merkel) beta images separately. The Yeo parcellation scheme was chosen because 1) it is based on functional divisions determined in a large sample (1,000 participants) and 2) includes default network regions that are anatomically very similar to known divisions determined by task-based fMRI studies investigating the neural correlates of social cognition, including self-reflection (Amodio & Frith, 2006; Denny et al., 2012; Saxe & Kanwisher, 2003). We calculated a one sample t-test on the Fisher z-transformed correlation values, in order to assess which ROIs yielded a correlation consistently above 0. Thresholded maps were generated using a false discovery rate (FDR) of 0.05 to correct for multiple comparisons.

### ROI-Based Pattern Similarity Analysis

To next assess whether voxels previously associated with the self demonstrate temporal compression, we employed an ROI approach using a Neurosynth-derived (Yarkoni et al., 2011) MPFC mask using the search term ‘self.’ This ROI has been used in prior work on self-representation (Denny et al., 2012; Lieberman et al., 2019; Meyer & Lieberman, 2018; Courtney & Meyer, 2020). Voxels in the association map test were restricted to Brodmann Area 10, a region particularly associated with self-representation (Denny et al., 2012; Lieberman et al., 2019; Meyer & Lieberman, 2018). Consequently, the MPFC ROI was restricted from −18 to 18 in the x dimension, 30 to 80 in the y dimension, and −12 to 22 in the z dimension, for a final mask size of 404 voxels (see Figure 4a).

For every contiguous time point for both the self and other (e.g., Self-Today and Self-3 Months in the Future), similarity between the two masked beta images was calculated via Spearman rank correlations within each subject. We then constructed a linear mixed model with target, the logarithmic time, and the interaction between target and logarithmic time as fixed effects predicting similarity (R values). We also included random intercepts and slopes for participants. MPFC ROI data and code have been made available on OSF (https://osf.io/m458t/).

## Acknowledgements

We would like to thank Matthew Lieberman, Naomi Eisenberger, and Mark Thornton for their feedback on this manuscript and Frances Sperry for her assistance with data collection.

The authors declare no competing interest.

---

**Supplementary Table 1.** Brain regions associated with the logarithmic model indicating past and future selves are collectively represented and compressed with time away from the present. Statistical significance was determined by a false discovery rate (FDR) of 0.05 to correct for multiple comparisons.

| Network | Region | x | y | z | p-value |
| --- | --- | --- | --- | --- | --- |
| **Core Default Network** | Medial Prefrontal Cortex | 0.6 | 50 | 5.2 | 0.003 |
|  | Posterior Cingulate | -0.6 | -50.8 | 31 | 0.0002 |
|  | Middle Temporal Gyrus | 61.8 | -6.8 | -18.2 | 0.008 |
|  | Temporoparietal Junction | 51.4 | -57.4 | 29.6 | 0.006 |
|  | Superior Frontal Gyrus | -22.4 | 28.8 | 46.6 | 0.002 |
| **MTL Default Network** | L. Precuneus | -12.4 | -55.4 | 12.6 | 0.005 |
|  | R. Precuneus | 13.8 | -53.2 | 13.8 | 0.001 |
|  | Middle Occipital Lobule | -39 | -80.4 | 32 | 0.005 |
| **DMPFC Default Network** | Angular Gyrus | -52.4 | -55.6 | 28.6 | 0.004 |
| **Frontoparietal Control A Network** | Middle Frontal Gyrus | -28.4 | 13.2 | 59.8 | 0.003 |
|  | Inferior Parietal Lobule | -48.8 | -53 | 49.2 | 0.003 |
| **Frontoparietal Control B Network** | Inferior Frontal Gyrus | -43 | 22.2 | 23 | 0.005 |
|  | Temporoparietal Junction | 40.8 | -49.6 | 45 | 0.005 |
| **Frontoparietal Control C Network** | Precuneus | 1.8 | -62.2 | 44.8 | 0.0008 |
| **Dorsal Attention A Network** | R. Middle Temporal Gyrus | 36.4 | -61.6 | 26 | 0.002 |
|  | L. Middle Temporal Gyrus | -33.6 | -65.8 | 23 | 0.0004 |
| **Visual A Network** | Primary Visual Cortex | 1.6 | -84.4 | -2.4 | 0.0002 |
| **Visual B Network** | Primary Visual Cortex | 1.6 | -71.4 | 11.6 | 0.0002 |

**Supplementary Table 2.** Demographic information for behavioral Mturk Studies 1-3.

| MTurk Studies | Gender (Women) | Age | Racial Breakdown; N (%) |  |  |  |  |  |  |
| --- | --- | --- | --- | --- | --- | --- | --- | --- | --- |
|  | N (%) | Mean (SD) | White | African American | Asian | Hispanic | Native American | Mixed Race | Other |
| **Study 1** | 85 (43%) | 35.9 (11.2) | 144 (73.5%) | 20 (10.2%) | 11 (5.6%) | 11 (6.4%) | 7 (3.6%) | 3 (1.5%) | - |
| **Study 2** | 91 (48.9%) | 35.2 (11.0) | 133 (71.5%) | 24 (12.9%) | 8 (4.3%) | 12 (6.5%) | 2 (1.1%) | 4 (2.2%) | 3 (1.6%) |
| **Study 3** | 95 (47.0%) | 35.9 (11.0) | 148 (73.2%) | 20 (9.9%) | 11 (5.4%) | 13 (6.4%) | 1 (0.5%) | 6 (3.0%) | 2 (1.0%) |

